# A systematic review and meta-analysis of the effects of older age on skeletal muscle mitochondrial function, as measured by ^31^P magnetic resonance

**DOI:** 10.64898/2026.05.02.722217

**Authors:** Donnie Cameron, Allan Clark, Laura J. Vermeulen, Arjan Malekzadeh, Vassilios S. Vassiliou, Melissa T. Hooijmans

## Abstract

**Objective:** Loss of skeletal muscle mass and performance is a hallmark of ageing. Mitochondrial function has been suggested as a critical determinant of skeletal muscle performance. However, mixed results have been reported regarding mitochondrial function in older individuals. Therefore, the primary objective of this systematic review is to determine whether ^31^P-MRS-derived τ_PCr_, reflecting mitochondrial oxidative capacity, is reduced in ageing skeletal muscle.

**Methods:** A preregistered systematic literature review was performed using the databases MEDLINE, EMBASE, SPORTDiscus, and Cochrane Central Register of Controlled Trials (CENTRAL). Papers were included if they reported τ_PCr_ as measured by ^31^P-MRS; and studied individuals over 65 years of age in combination with a younger control group. Differences between young and older groups were assessed using random effects meta-analysis.

**Results:** We included 20 papers in total, of which 2 measured 2 muscles, 5 focused on the tibialis anterior (TA) muscle, 11 on the calf muscles, 5 on the quadriceps, and 1 on the flexor digitorum longus. No statistically-significant differences were found in τ_PCr_ between older and younger adults for all muscles combined (Hedges’ *g*=0.11 (*p*=0.487). Inter-study heterogeneity was high (τ^2^=0.36, *I*^2^=72.49%, *H*^2^=3.64). Sub-analyses for the individual muscles showed longer τ_PCr_ in the quadriceps (*g*=0.65, *p*<0.001) in older adults, but shorter τ_PCr_ in the TA muscle (*g*=−0.64, *p*<0.001). For the calf muscles, no differences were detected between older and young individuals (*g*=0.20, *p*=0.377).

**Conclusion:** No uniform age-related decline was found for τ_PCr_ when comparing all studies together. Substantial heterogeneity was observed between the individual muscles, with τ_PCr_ being prolonged in the upper leg muscles in older adults, but shortened in the tibialis anterior. This suggests more work using standardised settings and well-defined cohorts is needed.

## INTRODUCTION

Ageing is closely associated with progressive declines in skeletal muscle mass and strength, which significantly impact mobility, independence, and quality of life in older adults^1–3^. This decline not only affects individuals but also places an increasing burden on healthcare systems, driving up injury rates, morbidity, and long-term care costs^4,5^. Beyond the physical consequences, a fear of falling can lead to reduced physical activity and social engagement, and loss of independence, contributing to further deconditioning and diminished well-being^6,7^. Despite its impact, the underlying mechanisms driving age-related declines in physical performance remain incompletely understood. A key determinant of physical performance is skeletal muscle mitochondrial function; however, mixed findings have been reported regarding how this is affected by ageing.

Phosphorus-31 magnetic resonance spectroscopy (^31^P-MRS) has emerged as a powerful non-invasive tool for evaluating mitochondrial oxidative capacity *in vivo*^8,9^ enabling the assessment of adenosine triphosphate (ATP), phosphocreatine (PCr), and inorganic phosphate (Pi) at rest and during, or after, voluntary exercise or electrical stimulation (Figure 1). Where resting metabolites measured with ^31^P-MRS provide information about tissue pH and basal ATP synthesis rate, exercise studies provide valuable information on mitochondrial oxidative capacity, which is proportional to the depletion of PCr during exercise, and its subsequent recovery. Indeed, the time constant of phosphocreatine recovery (τ_PCr_) after exercise is considered a direct measure of mitochondrial oxidative capacity, provided that oxygen delivery by the vascular system is not limiting^10^. A prolonged τ_PCr_ suggests impaired mitochondrial function, making it a valuable metric for comparing age-related differences in oxidative capacity. Other measures derived from ^31^P-MRS are: the initial rate of PCr recovery (Vi_PCR_) directly after exercise cessation, indicating how quickly the mitochondria start rebuilding PCr; and the maximal rate of oxidative ATP synthesis (Q_max_), which reflects the theoretical maximal mitochondrial ATP production, with higher values indicating greater recovery capacity. Both measures are derived using calculations based on different models and assumptions^9^. While ^31^P-MRS is valuable for studying muscle metabolism, findings on τ_PCr_ changes in ageing skeletal muscle are mixed.

**Figure 1.**
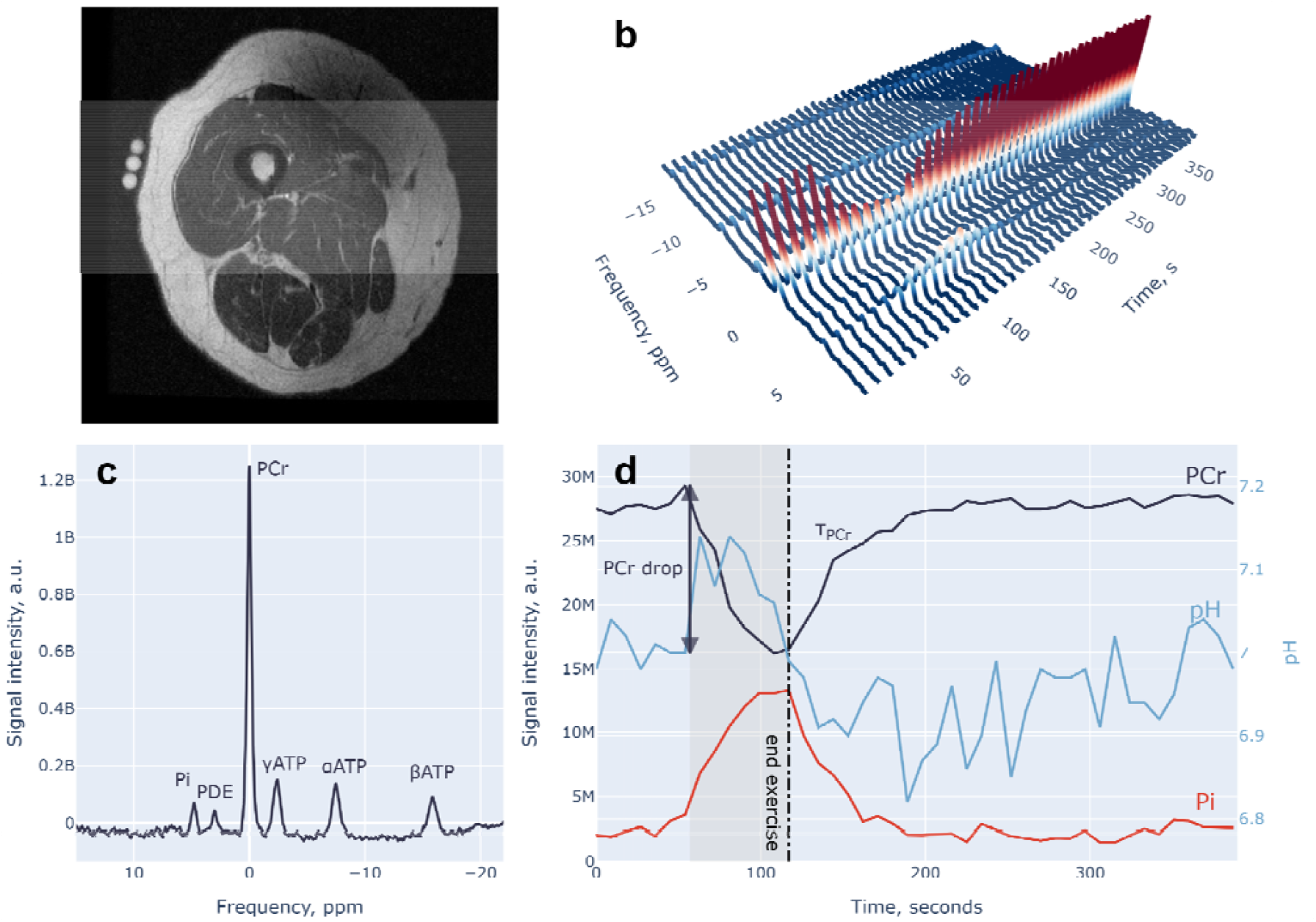
Example data from an exercise phosphorus-31 magnetic resonance spectroscopy (^31^P-MRS) experiment: a) verification of ^31^P coil positioning (centred on three fiducial markers) on an axial view of the upper leg; b) stacked ^31^P spectra obtained during rest, voluntary exercise, and recovery, showing the dynamics of phosphocreatine (PCr) and inorganic phosphate (Pi); c) a rest ^31^P spectrum showing other key metabolites—including phosphodiesters (PDE) and adenosine triphosphate (ATP); and d) a simplified representation of b) showing concentrations of PCr and Pi, and intracellular pH, during the course of the protocol, along with the PCr recovery time constant, τ_PCr_. An interactive version of this figure is available online (https://dc-3t.github.io/p31_sreview)

**Figure 2.**
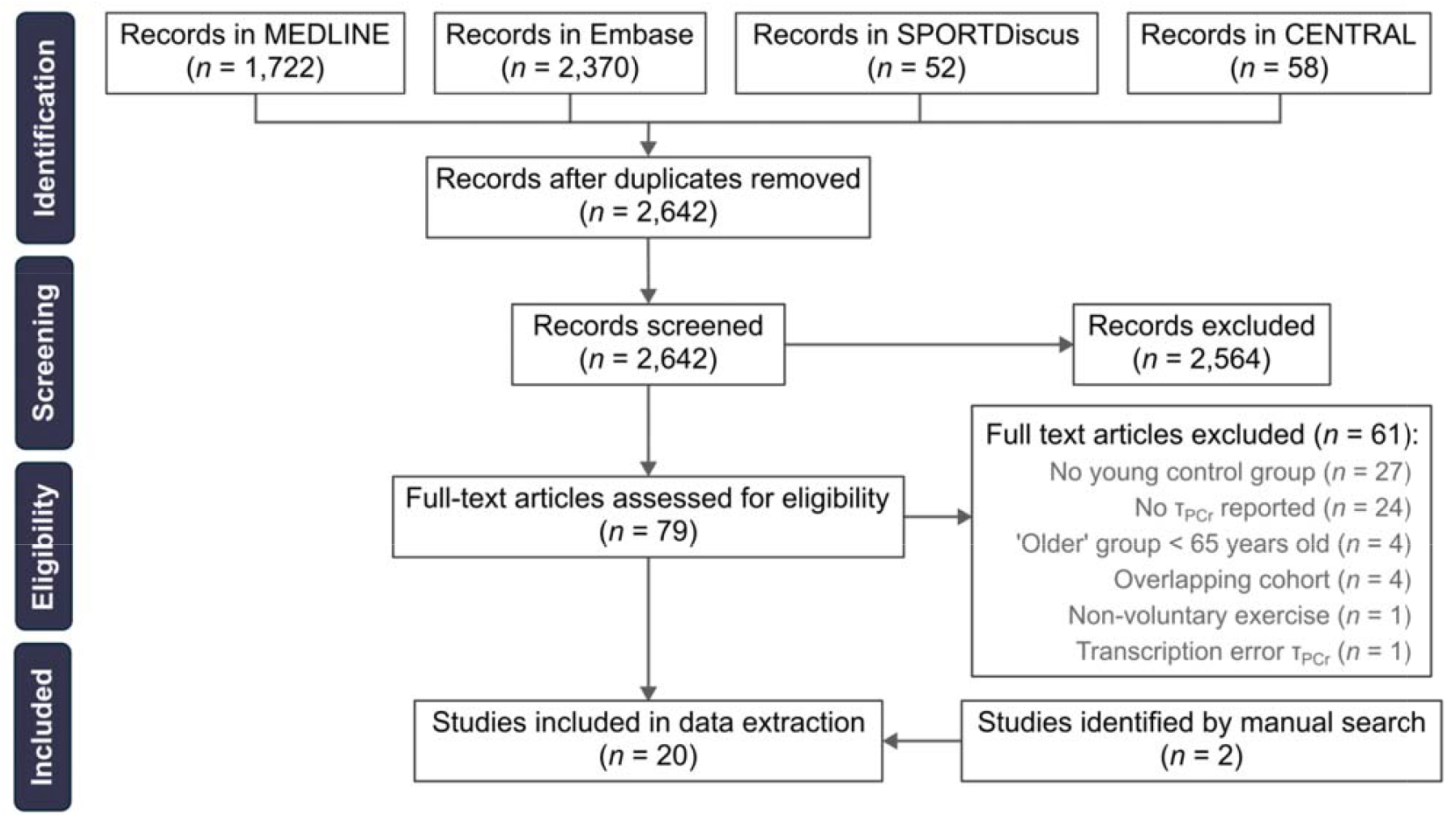
PRISMA flow diagram showing papers identified, screened, excluded, and ultimately included in this systematic review and meta-analysis

Given the essential role of mitochondrial oxidative capacity in skeletal muscle function, it is crucial to establish whether this parameter declines with age and is therefore linked to functional decline in older adults. The primary objective of this systematic review is to determine whether ^31^P-MRS-derived τ_PCr_, reflecting mitochondrial oxidative capacity, is reduced in ageing skeletal muscle. We also conduct sub-analyses by muscle group, physical activity control, and exercise protocol (dynamic versus isometric), and assess age-related changes in other metabolic measures, including end-exercise Pi, Vi_PCr_, and Q_max_. By addressing these objectives, we aim to clarify the relationship between ageing and mitochondrial oxidative capacity, providing a foundation for targeted interventions to maintain muscle health and quality of life in ageing populations.

## METHODS

### Search strategy and selection criteria

This systematic review was guided by the standards of the Preferred Reporting Items for Systematic Reviews and Meta-Analyses (PRISMA) statement^11^. The study protocol was registered in the International Prospective Register of Systematic Reviews (PROSPERO; registration number CRD42024494732). We searched the MEDLINE, EMBASE, SPORTDiscus, and Cochrane Central Register of Controlled Trials (CENTRAL) databases, from inception to the 10th of February, 2026. Preprints and conference abstracts were excluded. The search strategy included terms related to ^31^P-MRS, skeletal muscle, ageing, and mitochondrial function, with filters to exclude animal and oncological studies and review articles (see *Supplementary Methods S1*). Search results were imported into Rayyan (www.rayyan.ai) for independent abstract screening by two authors (D.C., M.T.H.). Duplicates were removed using a locally-developed tool (http://dedupendnote.nl:9777, version 1.06) and irrelevant studies were excluded based on predetermined inclusion and exclusion criteria. Studies were included if they: reported τ_PCr_ measured by ^31^P-MRS; and studied older individuals aged 65 years or older in combination with a younger control group. Studies using non-voluntary (electrically-stimulated) exercise were excluded. Where cohorts were reported in more than one study, with the same outcomes, only the study with the largest sample size was included.

### Data extraction

Relevant data from the included studies were independently extracted by two authors (D.C., M.T.H.), who resolved any disagreements by discussion. Extracted data included: per cent PCr depletion; τ_PCr_; end-exercise Pi/PCr, [PCr], [Pi], and pH; Pi recovery time; Vi_PCr_; Q_max_; Pi/PCr slope; and Pi/PCr and pH intracellular thresholds. Where K_PCr_ or PCr half-time (t_1/2_) were reported, these were converted to τ_PCr_ using the equations τ_PCr_ = 1/K_PCr_ and τ_PCr_ = t_1/2_/ln(2), respectively. Where data were missing or appeared to include errors, we contacted the corresponding author of the relevant papers. If data were presented graphically rather than numerically, we used the online graphreader tool (www.graphreader.com; website used in 2024) to extract mean values or individual data points, where appropriate. Further, when results were split by gender, we performed weighted averaging to determine overall mean values for older and younger groups. When studies included multiple younger and older groups, we preferentially included inactive-to-moderately-active individuals over athletic or frail individuals, or groups with explicitly comorbid conditions. Likewise, when results from multiple contraction protocols were presented, we included only those with short, fixed-weight bouts over steady-state or ramped protocols.

### Quality assessment

Study quality was assessed independently by two raters (D.C., M.T.H) using a version of the Newcastle–Ottawa Scale (NOS) adapted for cross-sectional studies (*Supplementary Methods S2*). Each study was awarded up to eleven stars, with higher scores indicating lower risk of bias. Discrepancies between the reviewers were resolved by discussion.

### Statistical analysis

All statistical analyses were performed in STATA (version 19.5, STATA Corp, College Station, TX). For each outcome, a random effects meta-analysis was performed using restricted maximum likelihood (REML) estimation. Given the small number of studies on this topic, effect sizes were calculated as Hedges’ g standardised mean differences with 95% confidence intervals, where positive values indicated a longer τ_PCr_ in older muscles. Effect sizes were interpreted as small (0.2), medium (0.5), and large (0.8)^12^. Heterogeneity was assessed via the *I*^2^ index (with 25%, 50%, and 75% indicating low, medium, and high heterogeneity, respectively) and Cochrane’s *Q* statistic. Statistical significance was set at *p*<0.05. In anticipation of substantial between-study heterogeneity, we performed subgroup meta-analyses of τ_PCr_ for different muscles or muscle groups and for different contraction protocols. A further meta-analysis of τ_PCr_ was performed after excluding studies that did not control for physical activity.

Publication bias was visualised using a funnel plot of effect size versus standard error and was quantitatively assessed via the Egger regression test. The symmetry of the funnel plot was visually inspected to identify potential small-study effects.

## RESULTS

### Search results

The literature search resulted in a total of 2,642 records (Figure 1). After screening titles, abstracts, and full texts according to the eligibility criteria, 18 studies were included. A manual search of the reference lists of these records yielded a further 2 studies, giving a total of 20 (*Supplementary Table S1*).

### Study characteristics

From the 20 studies that reported τ_PCr_ in young and older adults across different muscle groups^13–32^, 22 comparisons were possible, as two studies reported data from more than one muscle group. In total: five comparisons focused on the upper leg muscles, of which two assessed the quadriceps and three the vastus lateralis muscle; eleven investigated the calf muscles, with four focusing on the gastrocnemius medialis and one assessing both heads of the gastrocnemius; five examined the tibialis anterior; and one examined the flexor digitorum superficialis (FDS), in the forearm. Regarding secondary outcomes: six studies included end-exercise Pi, four included V_PCr_, and ten included Q_max_.

Important study characteristics are reported in *Supplementary Table S1*. A mix of MRI scanner vendors, field strengths, and coil hardware were used, whereas acquisition protocols mostly consisted of a pulse-acquire sequence with repetition times ranging from 2 to 4 seconds. The physical activity levels of participants varied across the included studies (sedentary to moderately active), and were not reported in some cases. Both dynamic exercise and isometric contraction protocols were used, with varying intensities and durations.

### Quality Assessment

Across the included studies, NOS scores ranged between 1 and 8 stars (median = 6; *Supplementary Results S1*), indicating middling methodological quality overall. Most studies scored well on comparability and suitability of outcomes, whereas selection of representative participants was less consistently addressed. Inspection of the funnel plot (Figure S1) suggested possible asymmetry indicative of small-study effects, with Egger’s test giving *p* = 0.07. This may indicate under-reporting of non-significant results.

### Phosphocreatine recovery time constant (τ_PCr_)

Examining τ_PCr_ between older and younger adults across various muscle groups (Figure 3), the overall effect size was 0.11 (*p* = 0.487), indicating no difference. Heterogeneity was high (τ^2^ = 0.36, *I*^2^ = 72.49%, *H*^2^=3.64; *Q* = 74.43, *p* < 0.001), implying substantial variability in the reported effect sizes.

**Figure 3.**
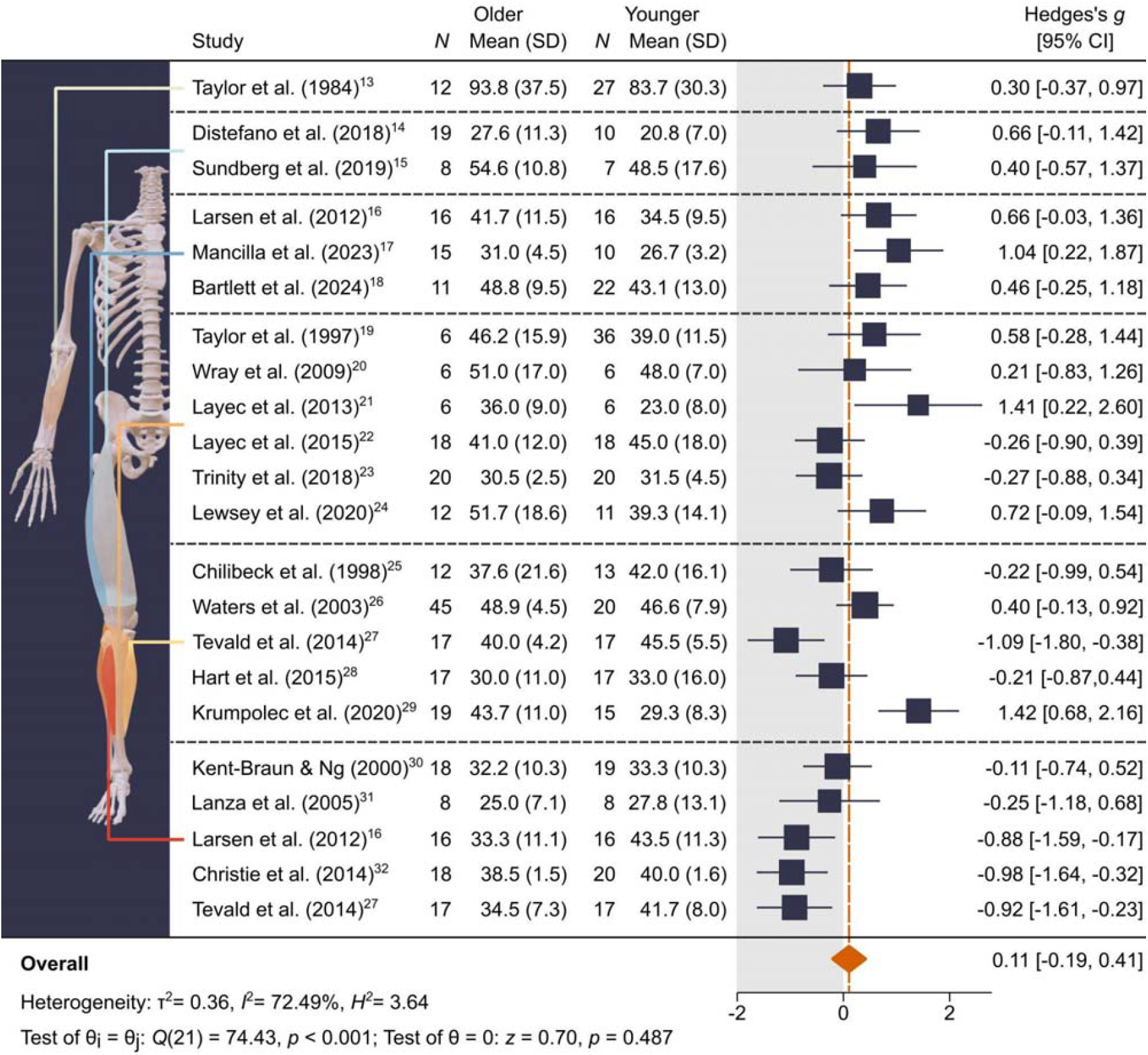
Forest plot of the difference in τPCr between older and younger adults based on a random effects meta-analysis model fit using restricted maximum likelihood. Results are shown for different muscles and muscle groups, arranged from superior (top) to inferior (bottom): flexor digitorum superficialis; quadriceps muscles; vastus lateralis; whole calf (triceps surae); gastrocnemius medialis; and tibialis anterior. Anatomy visualisations generated using AnatomyTOOL (https://anatomytool.org/open3dmodel)

#### Subgroup meta-analyses of individual muscles/muscle groups showed mixed results

The overall effect size was 0.53 (*p* = 0.009) for muscles in the anterior compartment of the upper leg (quadriceps and vastus lateralis), with older adults exhibiting significantly longer τ_PCr_ values (Figure 4). Heterogeneity was low (τ^2^ < 0.01, *I*^2^ < 0.01%, *H*^2^ = 1.00; *Q* = 1.39, *p* = 0.846), suggesting that inter-study differences were relatively minor.

**Figure 4.**
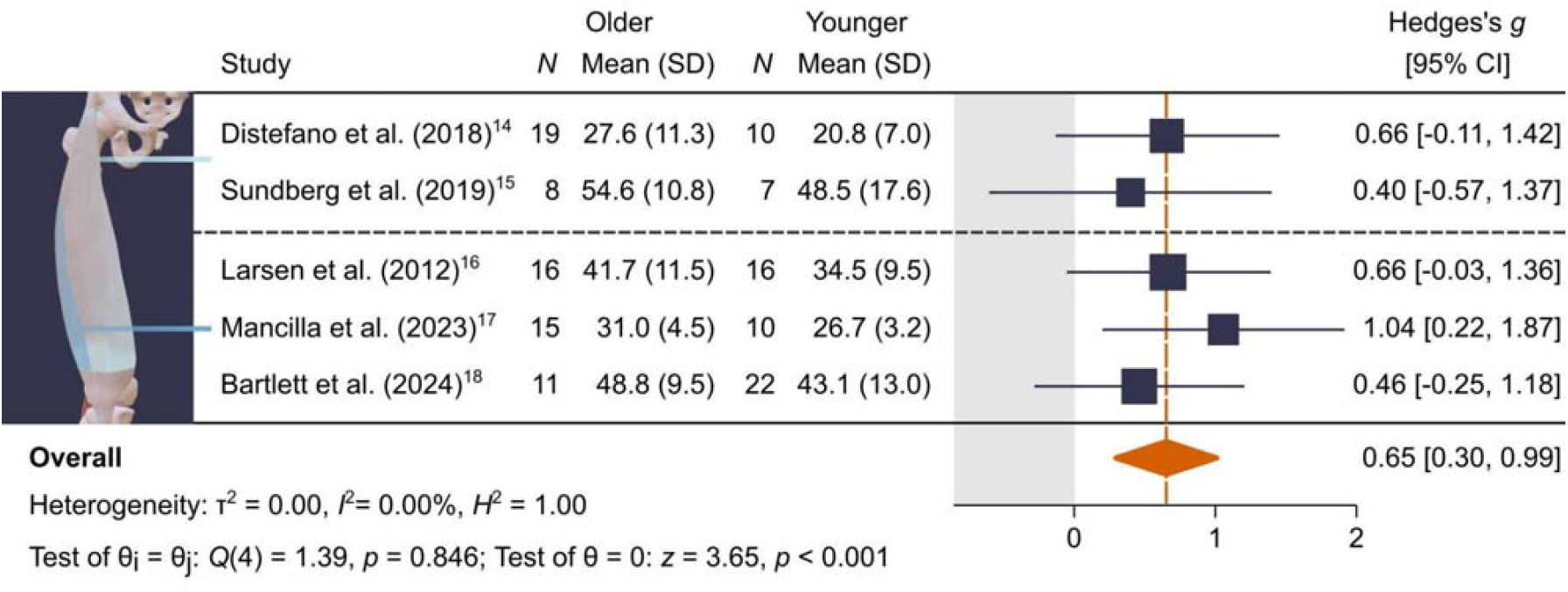
Forest plot of the difference in τPCr in muscles of the anterior compartment of the upper leg (quadriceps or vastus lateralis) between older and younger adults. Results are based on a random effects meta-analysis model fit using restricted maximum likelihood. Anatomy visualisations generated using AnatomyTOOL (https://anatomytool.org/open3dmodel)

In the calf and gastrocnemius medialis muscles (Figure 5a), the overall effect size was 0.20 (*p* = 0.377), indicating a nonsignificant difference between age groups. Effect sizes varied widely across studies, with some reporting prolonged τ_PCr_ in older adults and others reporting the opposite. Substantial heterogeneity was observed (τ^2^= 0.40, *I*^2^ = 74.09%, *H*^2^ = 3.86; *Q* = 36.56, *p* < 0.001), suggesting that inter-study differences may have influenced results.

**Figure 5.**
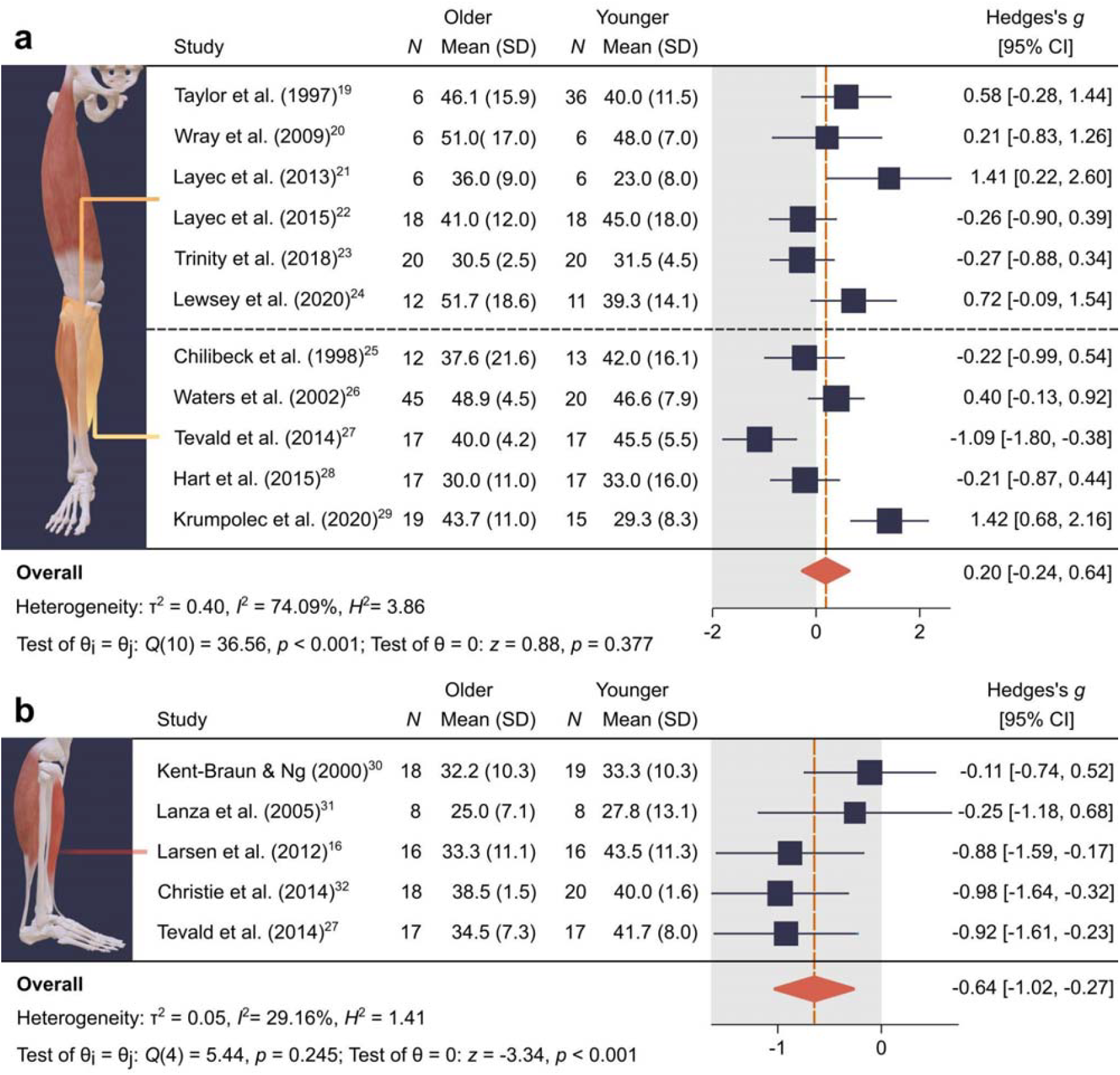
Forest plot of the differences in τPCr in muscles of the posterior compartment of the lower leg (panel a, whole calf or gastrocnemius medialis) and the tibialis anterior muscle (panel b) between older and younger adults. Results are based on a random effects meta-analysis model fit using restricted maximum likelihood. Anatomy visualisations generated via AnatomyTOOL (https://anatomytool.org/open3dmodel)

For the tibialis anterior (Figure 5b), the effect size was −0.64 (*p* < 0.001), indicating significantly shorter τ_PCr_ values in older adults. Heterogeneity across studies was relatively low (τ^2^ = 0.05, *I*^2^ = 29.16%, *H*^2^=2.98; *Q* = 5.44, *p* = 0.245).

#### No age differences relating to contraction protocol, physical activity, or secondary outcomes

When evaluating dynamic (effect size, *g* = 0.27, *p* = 0.075) and isometric (*g* = −0.11, *p* = 0.68) exercise separately, and including only studies that corrected for physical activity (*g* = −0.01 *p* = 0.94) no differences in τ_PCr_ were found between older and younger adults (*Supplementary Figures S2-S3*). Heterogeneity between studies was moderate to high in all these sub-analyses. Further, no differences in end-exercise Pi concentrations (effect size, *g* = 0.21; *p* = 0.48), Vi_PCr_ (g = 0.20 p = 0.62) or Q_max_ (g = 0.072; *p* = 0.77) were found between older and younger adults (Supplemental Figures S4-S5).

## DISCUSSION

This systematic review aimed to identify differences in skeletal muscle oxidative capacity, measured with ^31^P-MRS, between younger and older adults, with the expectation that older individuals have a prolonged τ_PCr_ recovery. When including all studies and muscle groups, no significant difference in τ_PCr_ was found; however, significant heterogeneity was observed between studies, suggesting that differences between muscle type, methodology, and participant characteristics may have influenced the results. This prompted sub-analyses across muscle groups, where we observed contrasting relationships in the quadriceps and tibialis anterior muscles: prolonged or shortened τ_PCr_, respectively, in older versus younger adults.

### Differences in τ_PCr_ in individual muscles or muscle groups

Analyses by muscle group revealed distinct age-related patterns in τ_PCr_. Muscles in the anterior upper leg consistently showed prolonged τ_PCr_ in older adults, with low heterogeneity across studies, suggesting a robust age-effect. The posterior compartment of the lower leg showed no significant age difference and considerable heterogeneity between studies, whereas TA studies reported a significantly shorter τ_PCr_ in older adults, suggesting preserved or even enhanced recovery dynamics. This difference, compared to other muscles, could be attributed to the TA muscle’s distinct fibre composition (Type I ~70-84%) and the selective Type II fibre atrophy associated with ageing^33,34^. Some studies report increases in Type I fibre proportion (up to 73–84%) in muscles of older individuals^35^, which may partly explain lower τPCr values in certain muscles. In theory, this shift should enhance oxidative capacity in Type II dominant muscles, yet the vastus lateralis still shows a pronounced age-related decline in τPCr. This apparent paradox likely reflects a decline in mitochondrial quality, rather than quantity, with age, potentially due to impaired mitophagy or increased oxidative damage, which can limit oxidative function even in Type I-enriched muscles^36^. Alternatively, there have been some reports that mitochondrial respiratory function is only reduced in glycolytic fibres^37–39^, suggesting another possible explanation. Beyond fibre types, heterogeneous muscle involvement may arise from age-related differences in daily activity, with more pronounced atrophy in muscles typically used for high-intensity, explosive movements. Finally, other ^31^P-MRS-derived metrics—end-exercise Pi, V_PCr_, and Q_max_—showed no consistent age differences, overall or per muscle/muscle group. Both V_PCr_ and Q_max_ showed considerable heterogeneity, whereas end-exercise Pi appeared homogeneous, contrasting with a recent systematic review that reported significantly higher end-exercise Pi in younger individuals^40^.

### The relationship between physical activity and τ_PCr_

In line with two recent “cross-talk letters” highlighting the importance of physical activity in ageing research, it is evident that disuse can produce muscle changes that closely mimic ageing^41,42^. Therefore, a key challenge in interpreting age-related differences in muscle oxidative capacity is the inconsistent reporting of participants’ habitual activity levels across studies. In this review, a subset of four studies strategically included groups of highly-active or relatively frail older individuals to directly investigate the effect of physical activity^16,17,24,32^. They showed that mitochondrial dysfunction is more pronounced in less-active individuals, whereas trained older adults are often comparable to younger controls, even if differences are not always statistically significant. Similar patterns are supported by ex vivo studies when comparing older active with older sedentary individuals^43–45^ and by evidence from younger and middle-aged subjects following periods of inactivity^46–48^, underscoring the central role of physical activity. Notably, even when total activity is matched, younger individuals typically spend more time in moderate-to-vigorous intensity exercise, whereas older adults tend to engage primarily in low-intensity activity^16,27^. This suggests that exercise intensity, and not just the volume of exercise, is critical for maintaining mitochondrial function. A related and still unresolved question is whether there exists a specific “dose” of physical activity capable of fully preserving mitochondrial function with advancing age. What is clear, however, is that training interventions consistently demonstrate that mitochondrial respiratory function can improve across all ages, even in previously sedentary individuals^49^. This is supported by in vivo, in vitro, and ex vivo assessments^44,49,50^.

### The role of convective and diffusive O_2_ delivery on τ_PCr_ recovery

The interpretation of ^31^P-MRS measures of oxidative capacity is fundamentally predicated on the assumption that convective oxygen delivery and oxygen diffusivity are not limiting factors. However, previous work indicates that even submaximal dynamic exercise can induce oxygen limitations in PCr kinetics in both trained and sedentary subjects^51,52^. Consequently, to disentangle exercise limitations on the level of the peripheral muscle, it is necessary to measure not only mitochondrial function, but also convective and diffusive oxygen delivery, simultaneously with ^31^P-MRS^53^. Most studies in this review did not account for these factors, though some employed multimodal approaches—including MRI, near-infrared spectroscopy, and ultrasound or manipulated inspired air conditions— yielding important insights. Two groups performed popliteal flow and tissue oxygenation measurements alongside ^31^P-MRS and found no differences in PCr recovery rates between young and older individuals with similar or slightly higher flow and oxygenation^23,28^. Layec et al. observed a complex response to inspired air conditions: τ_PCr_ remained unaffected in older individuals, while it tended to be prolonged in younger individuals under hyperoxic conditions—a phenomenon the authors attributed to potential oxygen-induced vasoconstriction in the young^21^. Another study used advanced interleaved MRI to assess metabolic and oxygenation dynamics simultaneously^20^, finding reduced blood flow in older adults, but no difference in PCr recovery times. Despite this, older individuals showed greater PCr depletion and higher Pi/PCr ratios at end-exercise, indicating increased metabolic stress, while pH remained similar between groups. That τ_PCr_ was preserved despite reduced oxygen delivery suggests a stronger metabolic drive, with elevated ADP and Pi levels maximally stimulating mitochondrial function. This implies that intrinsic mitochondrial capacity (Q_max_ or ADP sensitivity), rather than oxygen supply, may be the primary determinant of recovery kinetics under these conditions. Alternatively, it is also plausible that the exercise protocols were not sufficiently demanding to elicit oxygen delivery limitations, thereby masking potential age-related differences.

### Dynamic and isometric exercise protocols

The type and duration of exercise play a critical role in determining the energy systems recruited during exercise, as does oxygen availability. Adequate oxygen availability is essential for initiating PCr resynthesis, and influences both PCr recovery kinetics and the metabolic pathways engaged during exercise. We did not observe differences in τ_PCr_ across the contraction protocols examined; however, this finding contrasts with the study by Bartlett and colleagues, who directly compared dynamic and isometric contraction protocols in young and older individuals^18^. In their study, PCr recovery was approximately 20% slower in young participants following an isometric protocol compared to dynamic protocol, whereas this difference was less pronounced in older individuals. The differential effects of contraction protocols on PCr recovery kinetics between young and older individuals suggest that future studies should carefully consider and standardise contraction protocols, particularly when examining age-related differences in muscle oxidative capacity.

### Linking ^31^P-MRS to biopsy findings

Alongside^31^ P-MRS, *ex vivo* and *in vitro* methodologies are commonly employed to investigate mitochondrial content, function, and morphology, but findings remain mixed. Studies report: impaired mitochondrial respiratory function in permeabilised fibres^43,54,55^; preserved respiratory function in permeabilised fibres but not in isolated mitochondria^56^; or no age-related differences^57^. The limited investigations of mitochondrial morphology are similarly inconsistent, including reports of increased fragmentation and smaller mitochondria, enlarged subsarcolemmal mitochondria in mice, and reduced mitochondrial size and area with advancing age in humans^58,59^. Importantly, these mixed results broadly align with *in vivo* measurements, though muscle-specific differences are rarely addressed. Notably, the TA muscle, despite being straightforward to sample *in vivo*, has been largely overlooked in biopsy studies, which mostly focus on the vastus lateralis or calf muscles. Consequently, the apparent preservation or improvement of ^31^P-MRS-measured oxidative capacity in the TA muscle with age cannot yet be confirmed through direct biopsy-based assessments. Additionally, biopsy samples are usually collected from the mid–muscle belly, failing to capture the full proximodistal heterogeneity of skeletal muscle tissue. This limitation is particularly relevant because PCr recovery kinetics vary along the length of the TA in healthy individuals^60^, and recent calf muscle studies reported that distal and proximal muscle regions are affected early in older individuals^61^, suggesting that mid-belly biopsies may underestimate age-related mitochondrial changes.

### Limitations

A significant challenge for data interpretation in this work was inconsistent reporting of crucial components such as the degree of PCr depletion, end-exercise pH, exercise intensity, participant physical activity levels, gender, and specific details regarding the ^31^P-MRS coil dimensions, placement, and exercise type and protocol. Indeed, the exercise protocols used to investigate mitochondrial function in the different studies varied greatly: from short (16-24-seconds) maximal voluntary isometric contractions to long dynamic exercise bouts (40-50% work rate for 5 minutes). Adherence to expert recommendations for standardised reporting would greatly enhance comparability in future research^9^. Methodological disparities in ^31^P-MRS acquisition also introduced variability. For instance, coil size and location directly dictate the sampled muscle volume, potentially influencing results if different studies sampled different fibre type compositions or muscle compartments. Variations in temporal resolution (e.g., between two and ten seconds) could over- or underestimate changes in PCr and Vi_PCr_, indirectly influencing Q_max_, and potentially obscuring differences between groups. Additionally, calculations for Vi_PCr_ were sometimes not corrected for T_1_ relaxation effects, which can impact accuracy. Finally, several large cohort studies were excluded from this meta-analysis either because they lacked a control group or group comparison, or because their primary focus was not on assessing τ_PCr_^62–64^. Where possible, analyses of these datasets in line with the expert recommendations could shed further light on changes in mitochondrial function in ageing.

## CONCLUSION

This systematic review examined whether skeletal muscle oxidative capacity, as assessed by ^31^P-MRS, is reduced in older adults. Overall, τ_PCr_ did not show a consistent age-related decline. Instead, findings were heterogeneous: τ_PCr_ was prolonged in the upper leg muscles (quadriceps and vastus lateralis) but shortened in the tibialis anterior muscle in older adults. These discrepancies likely reflect muscle-specific physiology, participant variability, and methodological factors. More research is needed to clarify the underlying sources of heterogeneity in ^31^P-MRS-derived metrics and to evaluate whether they can reliably serve as indicators of mitochondrial function in ageing populations, particularly among sarcopenic or frail individuals. Standardised protocols, muscle-specific analyses, and inclusion of well-

characterised cohorts will be critical for advancing the field.

## SUPPLEMENTARY INFORMATION

Supplementary Table S1

Supplementary Figure S1

Supplementary Figure S2

Supplementary Figure S3

Supplementary Figure S4

Supplementary Figure S5

Supplementary Figure S6

Supplementary Methods S1

Supplementary Methods S2

Supplementary Results S1

