## Supplementary for "A systematic review and meta-analysis of the effects of older age on skeletal muscle mitochondrial function, as measured by ^31^P magnetic resonance"

**Supplementary tables**

**Table S1**

| **Author/year** | **Number of participants, Age & Muscle Investigated** | **Level of Physical Activity** | **Type of exercise** | **Level of PCr depletion (%)** | **Vendor & field strength/ coil/localisation & TR** |
| --- | --- | --- | --- | --- | --- |
| Layec 2013 | Young (N=6): 26(10) Older (N=6): 69(3)  **Men only**  Calf muscles | · Self-report and interview  · Sedentary individuals both young and old; no evidence of regular or occasional physical activity | · Dynamic plantarflexion exercise  · Intensity: 50% WR max  · Frequency: 1 Hz  · Duration: 4 minutes | · Not different between groups;  · Reported in mMol at end-exercise; 19 (young) versus 24 (old) mMol;  · Resting values not reported | · 1.5T GE system;  · Double tuned ^31^P (11.5 cm square) and ^1^H 2 14✕15.5 cm) coil from Medical Advances;  · Pulse acquire; TR: 4,000 ms |
| Layec 2015 | Young (N=18): 22(1.6) Older (N=18): 74(8.1)    Gender division ~ 50/50  Calf muscles | · Questionnaire, accelerometry data and step count  · No difference in physical activity between groups based on steps per day and accelerometer counts per minute | · Constant-load supramaximal plantar ﬂexion  · Intensity: 120% of WRmax  · Frequency: 1 Hz  · Duration: 1 minute | · Not different between the groups  · +/-60% PCr depletion for both groups | · 2.9T Siemens;  · Double tuned ^31^P-^1^H coil with linear polarisation (rapid) 125 mm (^31^p) and 110 (1h)  · Pulse-acquire; TR: 2,000 ms |
| Lewsey 2020 | Young (N=11): 50.5(2.1) Older (N=12): 78.8(2.0)    Gender division ~ 50/50    Calf muscles | · Not reported | · Plantar flexion exercise;  · Intensity: started with 0.9 kg, second stage add 0.9kg; third stage at 1.8kg.  · Frequency: 1 sec audio cues  · Duration: 6 min total; each stage was 120 seconds; | · Not different between groups  · Reported in mMOL recalculated; +/- 60 PCr depletion for both groups | · 3.0T Philips;  · Custom-built ^31^P-MRS coil;  · Pulse-Acquire; TR: 2,000 ms |
| Taylor 1997 | Range young (N=20): 20-29 Range older (N=6): 70-83    Gender division ~ 50/50    Calf muscles | · Not reported | · Dynamic Plantarflexion exercise  · Intensity: starting at 10% body mass; Incremental after 4 spectra with 2% lean body mass each additional spectrum  · Frequency of at 0.5 Hz  · Duration: 32 fids | · Not different between groups  · 52% (older) versus 58% (young) PCr depletion | · 2.0T oxford instruments magnet with Bruker spectrometer;  · 6 cm diameter loop ^31^P single tuned;  · Pulse-acquire; TR varies defined in FIDs post-exercise |
| Trinity 2018 | Young (N=20): 23(1) Older (N=20): 72(2);  Gender division ~ 50/50  Calf muscles | · Step count and accelerometry measurements  · Sedentary to moderately physically active;  · Matched young with older participants | · Submaximal plantarflexion exercise  · Intensity: 40% WR max  · Frequency: 1 Hz  · Duration: 5 minutes | · Not different between the groups;  · Reported in mMol difference -14,5 (young) versus -15Mmol (older). | · 3.0T Siemens;  · Double tuned ^31^P (125 mm) and ^1^H (110 mm) with linear polarisation  · Pulse-acquire; TR: 2,000 ms |
| Wray 2009 | Young (N=6): 26(5) Older (N=6): 70(5)  **Gender not reported**  Calf muscle coil is placed on the GCM | · IPAQ questionnaire  · All participants normally active (no regular exercise routine  · matched for activity | · Plantar flexion exercise  · Intensity: 5 W  · Frequency: 0.33 Hz  · Duration: 5 minutes | · Greater PCr depletion in older individuals;  · 28% (older) versus 19% (young) | · 4.0T Magnex 4/60;  · 17 cm ^1^H volume coil for imaging and an 8 cm ^31^P surface coil  · Interleaved with SATIR; TR: 3,000 ms |
| Chillibeck 1998 | Young (N=10): 27.5(2.0) Older (N=10): 66.9(3.7)  Gender division ~ 50/50  Lateral head of the Gastrocnemius | · Lifestyle ranged from sedentary to moderately active | · Dynamic ankle plantarflexion exercise;  · Intensity: 80% of subjects pH threshold  · Square wave plantar flexion tests | · 29.8% (young) versus 22.6% (older);  · not different between groups | · 1.5T Siemens;  · Dual tuned (^31^P/^1^H) surface coil (5 cm)  · Pulse acquire; TR: 1,000 ms |
| Hart 2014 | Young (N=20): 22.0(2.0) Older (N=20): 73.0(7.0);  Gender division ~ 50/50  GCM muscle | · Assessed using a structured interview and accelerometry  · Moderately active both groups;  · Matched for activity | · Submaximal dynamic plantarflexion exercise  · Intensity: 40% of WRmax  · Frequency: 1Hz  · Duration: 5 minutes | · Not different between groups;  · 40% (young) and 51% (older) PCr depletion | · 3.0T Siemens;  · Double tuned ^1^H/^31^P surface coil (110 mm/125 mm) loop with linear polarisation  · Pulse acquire; TR: 2,000 ms |
| Krumpolec 2020 | Young (N=15): 29.4(6.7) Older (N=19): 64.6(5.8);  **Gender not reported**  Medial head of the gastrocnemius | · Not reported | · Isometric plantar flexion exercise  · Intensity: 30% MVC;  · Frequency: every 2 seconds  · Duration: 6 min exercise | · Different between the groups;  · 23.1% (young) versus 47.9% (older) PCr depletion | · 7.0T Siemens;  · ^31^P/^1^H surface coil (10 cm Rapid Biomedical);  · DRESS localisation; TR: 2,000 ms |
| Tevald 2014 | Young (N=17): 24(3) Older (N=17): 69(3)  Gender division ~ 50/50  Medial head of the gastrocnemius | · Measured PA counts with accelerometry;  · Sedentary individuals  · not different in the LPA range;  · Different in the MVPA range | · plantarflexion isometric exercise  · Intensity: Maximal  · Duration:20 seconds | · Not different between groups  · 40.6% (older) /34.6%(young) | · 4.0T Bruker;  · ^31^P surface coil (elliptical 3✕5cm);  · Pulse acquire; TR: 2,000 ms |
| Waters 2003 | Young (N=20): 25(4) Older (N=45): 73(4)  Gender division ~ 50/50  GCM | · Assessed using a modified interview questionnaire  · Similar levels of Physical Activity | · Dynamic Plantar flexion exercise  · Frequency: Every 2 seconds  · IntenAt 15% of total lean body mass  · 12 minutes of exercise | · PCr depletion was more in the older individuals compared to the younger individuals  · 15% (young)/ 20% (older) | · 1.5T GE Healthcare;  · ^31^P transmit receive flex coil; (loop size not reported)  · Pulse-acquire; TR: 2,000ms & NSA ✕ 4 = actual TR: 8,000 ms |
| Kent-Braun & Ng 2000 | Young w (N=9): 33.2(4.6) Young m (N=10): 33.4(5.2) Older w (N=9): 75.3(4.6) Older m (N=9): 75.7(4.7)  Gender division ~ 50/50  TA muscle | · Measured with recall questionnaire and 3D accelerometry for 7 days  · No difference in physical activity measures; | · MVIC -ankle dorsiflexion exercise  · 15 seconds | · PCr depletion reported separately for men and woman & young woman 31.6%(4.9); older woman 33.1%(5.0); young men 33.9%(6.0) and older men 41.9%(5.4)  · Not different between young and older | · 1.9T; 30 cm bore Oxford magnet;  · ^31^P surface coil (elliptical 3✕5 cm);  · Pulse-acquire; 1,250; 2 NSA & 2,500 ms |
| Lanza 2005; | Young (N=8): 22(1) Older (N=8): 75(5)  **men only**    TA muscle | · Relatively sedentary  · Activity monitoring using accelerometry for 5 days  · not different between groups | · Ankle Dorsiflexion  · Intensity: maximal isometric  · Duration: 16 seconds and 60 seconds | · Not different between groups in 16 second condition; 20.7mMol/38.6mMol & 47% Depletion (young) and 22.4/37.3 & 40% Depletion  · Different and more depleted in the (80% depletion) young subjects/ 63% PCr depletion older subjects in 60 second condition. | · 4.0T Bruker;  · ^31^P surface coil (elliptical 3✕5 cm)  · Pulse-acquire; 2,000ms; averaged to have a 6sec temporal window |
| Larsen 2012 | Young m (N=8 ): 24.8(3.5) Young w (N= 8): 27.0(3.1) Older m (N=8): 68.9(3.9) Older w (N=8 ): 69.4(2.4)  Gender division: ~ 50/50  TA muscle | · Measured with accelerometry and questionnaires  · PA total time not different;  · Older spent more in LPA and young more MVPA | · Dorsiflexion exercise  · Intensity: maximal Isometric  · Duration: 16 seconds | · PCr Depletion not different between groups  · 51.7% (young) versus 60.5% (older) | · 3.0T Philips;  · ^31^P surface coil (elliptical 3✕4 cm)  · Pulse-acquire; TR: 2,000ms |
| Tevald 2014 | Young (N=17): 24(3) Older (N=17): 69(3)  Gender division ~ 50/50  TA muscle | · Measured PA counts with accelerometry;  · Sedentary individuals  · Not different in the LPA range;  · Different in the MVPA range | · Dorsiflexion exercise  · Intensity: maximal Isometric  · Duration: 16 seconds | · Not different between groups  · 38.7% (young) versus 42%(older) | · 4.0T Bruker;  · ^31^P surface coil (elliptical 3✕5 cm);  · Pulse-acquire; TR: 2,000ms |
| Christie 2014 | Young(N=20): 24(1) Older (N= 18 ): 73(2) Older impairments(N=9): 74(1)  Gender division ~ 50/50  TA muscle | · Accelerometry 10 day period;  · PA counts not different between young and older;  · PA counts different between older impaired and young and older | · 12 seconds contraction at 20% MVC, 50% MVC and at 100% MVC; | · Not different between groups  · Actual numbers not reported | · 4T Bruker;  · Double tuned ^1^H (7 cm circular) and ^31^P (3✕4 cm) surface coil;  · Pulse-acquire; TR: 2,000ms |
| Distefano 2018 | Young: 31.2(5.4) Older active (N=10):67.5(2.7) Older sed (N=19): 70.7(4.7)  Gender division ~ 50/50  Quadriceps muscles | · Accelerometry  · Young more active, in steps, Energy expenditure and active energy expenditure per 24h compared to older active and older sedentary | · Knee extension;  · Isometric kicking exercise; limited motion  · Duration: 45 seconds;  · Intensity: not reported  · Frequency: not reported | · Not reported | · 3.0T Philips;  · ^31^P surface coil (loop size not reported);  · Pulse-acquire; TR: 1,500ms; but averaged over 4 consecutive acquisitions with actual TR: 6,000ms |
| Sundberg 2019 | Young (N=7): 22.7 (1.2) Older (N=8): 76.4 (6.0)  Gender division ~ 50/50  Quadriceps muscles | · not reported | · Knee extension  · Intensity: maximal Isometric  · Duration: 24 seconds | · Not different between groups  · 52 & 56% PCr depletion young and older respectively; | · 3.0T GE Healthcare;  · ^31^P surface coil (12.7 cm square coil)  · Pulse-acquire; TR: 2,000 ms & averaged to actual TR: 8,000ms |
| Bartlett2024 | Young (N=22): 26.8(3.1) Older (N=11): 71.9(5.3)    Gender division ~ 50/50    Vastus Lateralis muscle | · Measured both PA count and MVPA using accelerometry measures  · similar activity levels between groups | · Knee extension exercise;  · Various contraction protocols;  · Intensity: Maximal Isometric  · Duration: 24 sec duration | · Not different between groups  · +/-50% PCr depletion | · 3.0T Siemens;  · Dual tuned ^31^P/^1^H (8✕10.5 cm);  · Pulse-acquire; TR: 2,000ms |
| Larsen 2012 | Young m (N=8 ): 24.8(3.5) Young w (N= 8): 27.0(3.1) Older m (N=8): 68.9(3.9) Older w (N=8 ): 69.4(2.4)  Gender division ~ 50/50  Vastus Lateralis muscle | · Measured with  · PA total time not different;  · Older spent more in LPA and young more MVPA | · Knee extension  · Intensity: maximal  · Frequency: Isometric  · Duration: 24 seconds | · No difference in PCr depletion between groups  · 50% (young) to 56% (older) PCr depletion; | · 4.0T Bruker;  · ^31^P surface coil (6×8 cm elliptical);  · Pulse-acquire; TR: 2,000ms |
| Mancilla 2023 | Young (N=10): 23(1) Older (N=15): 70(2)    Gender division ~ 50/50    Vastus Lateralis muscle | · Normally physically active maximum 1 structured exercise session per week and trained more than 3 exercise sessions of at least an hour;  · unclear how well matched | · Dynamic knee extension  · Intensity: 50-60% of the individuals predetermined maximum  · Duration: 5 minutes  · Frequency not reported | · Not reported | · 3.0T Philips;  · ^31^P surface coil (6 cm);  · Pulse-acquire; TR: 4,000ms |

**Table S1**. Characteristics of the included studies. ^1^H = proton, ^31^P = phosphorus-31, FID = free induction decay, LPA = light physical activity, MRS = magnetic resonance spectroscopy, MVC = maximum voluntary contraction, MVC = maximum voluntary isometric contraction, MVPA = moderate-to-vigorous physical activity, PA = physical activity, PCr = phosphocreatine, SATIR = saturation inversion recovery, TR = repetition time, WR = work rate

**Supplementary figures**

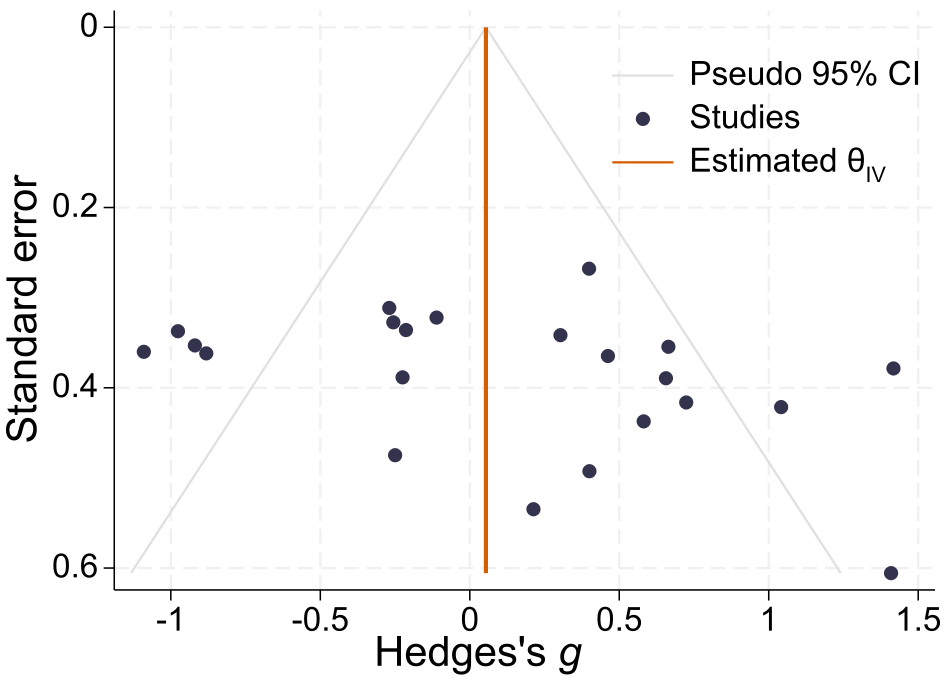

**Figure S1.** *Funnel plot showing the standard error of the effect-size estimate versus the Hedge’s* g *effect size of all studies included in this systematic review. Slight asymmetry may indicate under-reporting of non-significant results*

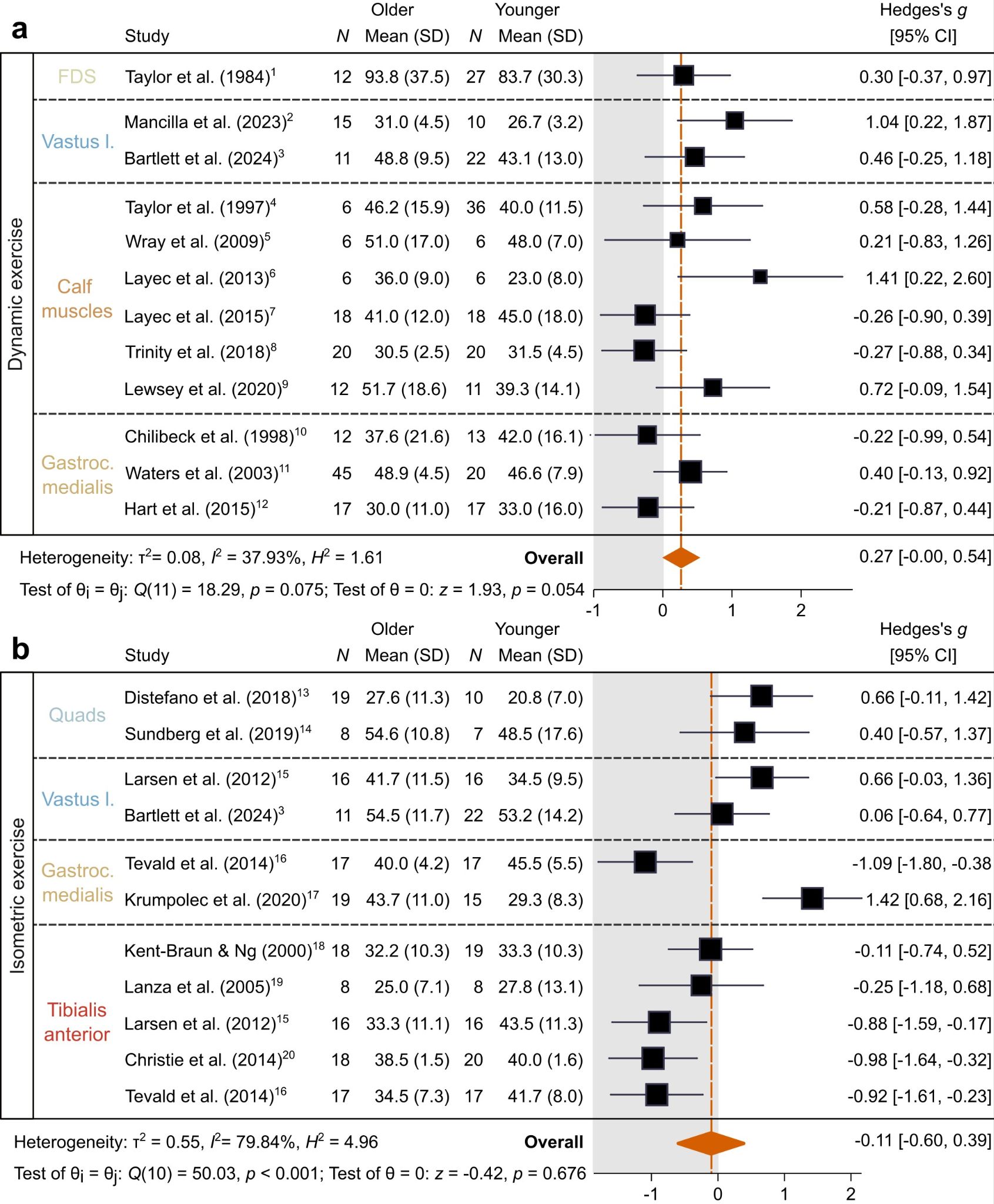

**Figure S2.** *Forest plot of the difference in τPCr between older and younger adults based on a random effects meta-analysis. Studies are divided into those that used a dynamic exercise protocol (panel a) or an isometric protocol (panel b). FDS = flexor digitorum superficialis, quads = quadriceps, vastus l. = vastus lateralis; gastroc. medialis = gastrocnemius medialis*

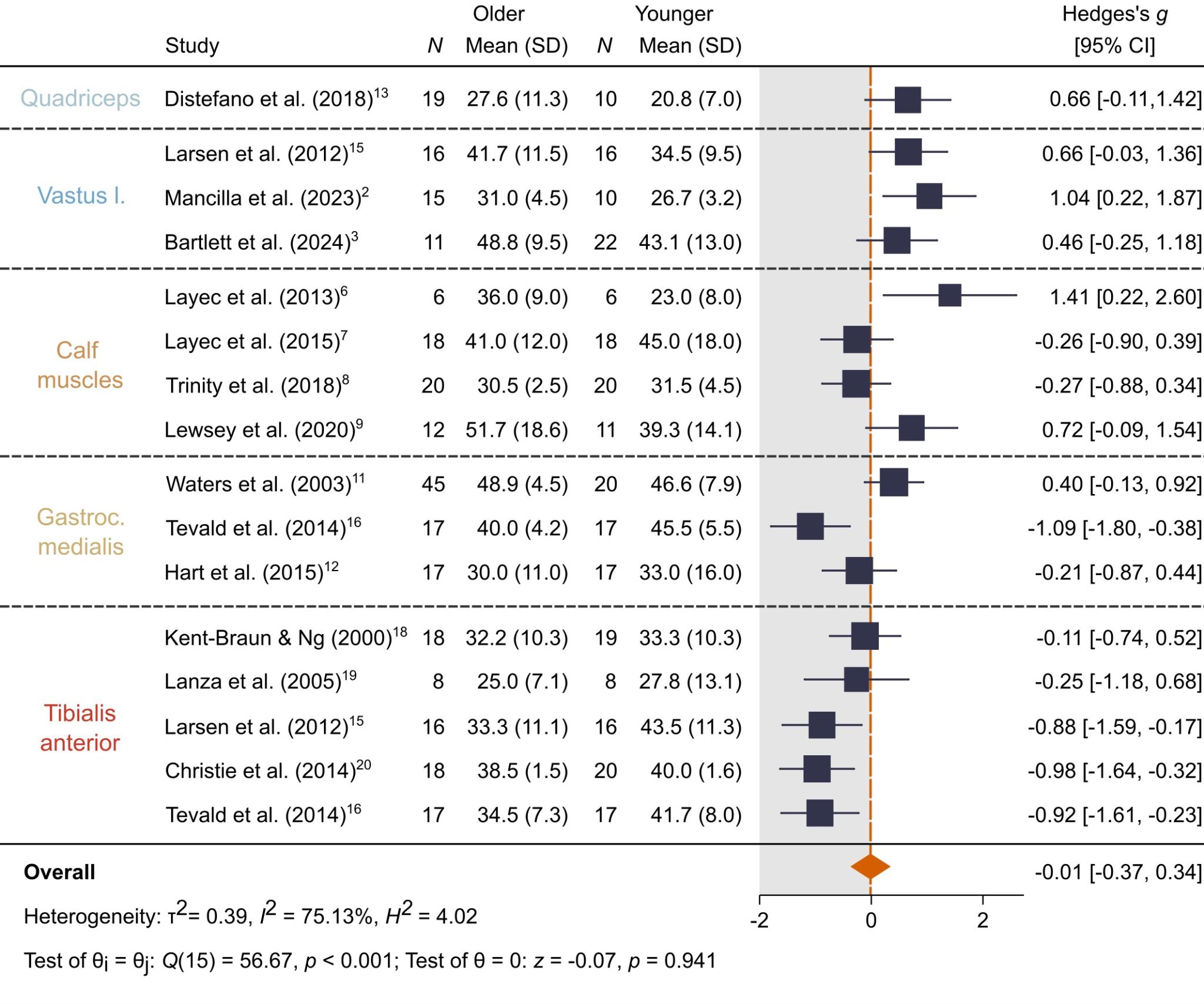

**Figure S3.** *Forest plot of the difference in τPCr between older and younger adults based on a random effects meta-analysis model fit using restricted maximum likelihood. Results are shown only for studies that controlled for physical activity levels, with different muscles and muscle groups arranged from superior (top) to inferior (bottom). Vastus l. = vastus lateralis; gastroc. medialis = gastrocnemius medialis*

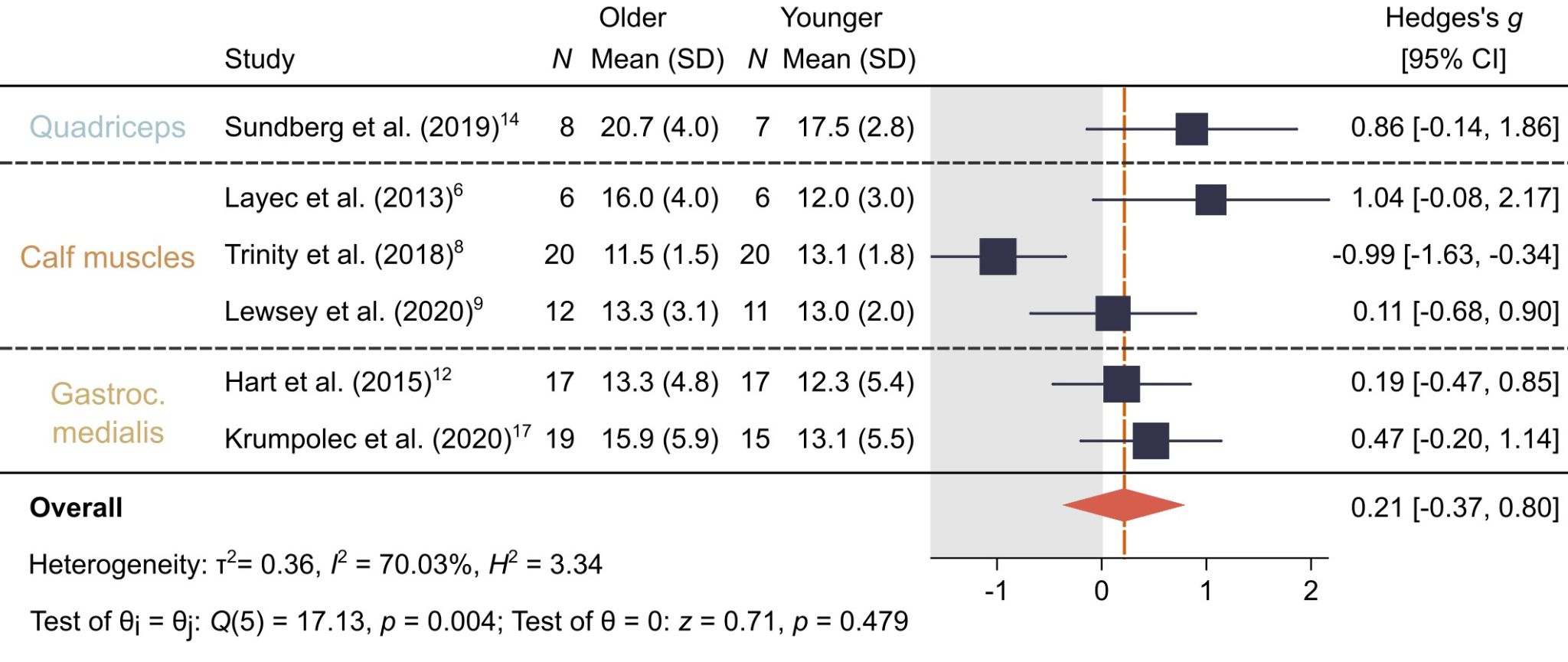

**Figure S4.** *Forest plot of the difference in end-exercise inorganic phosphate (Pi) concentrations in muscles of the upper and lower leg between older and younger adults. Results are based on a random effects meta-analysis model fit using restricted maximum likelihood. Gastroc. medialis = gastrocnemius medialis*

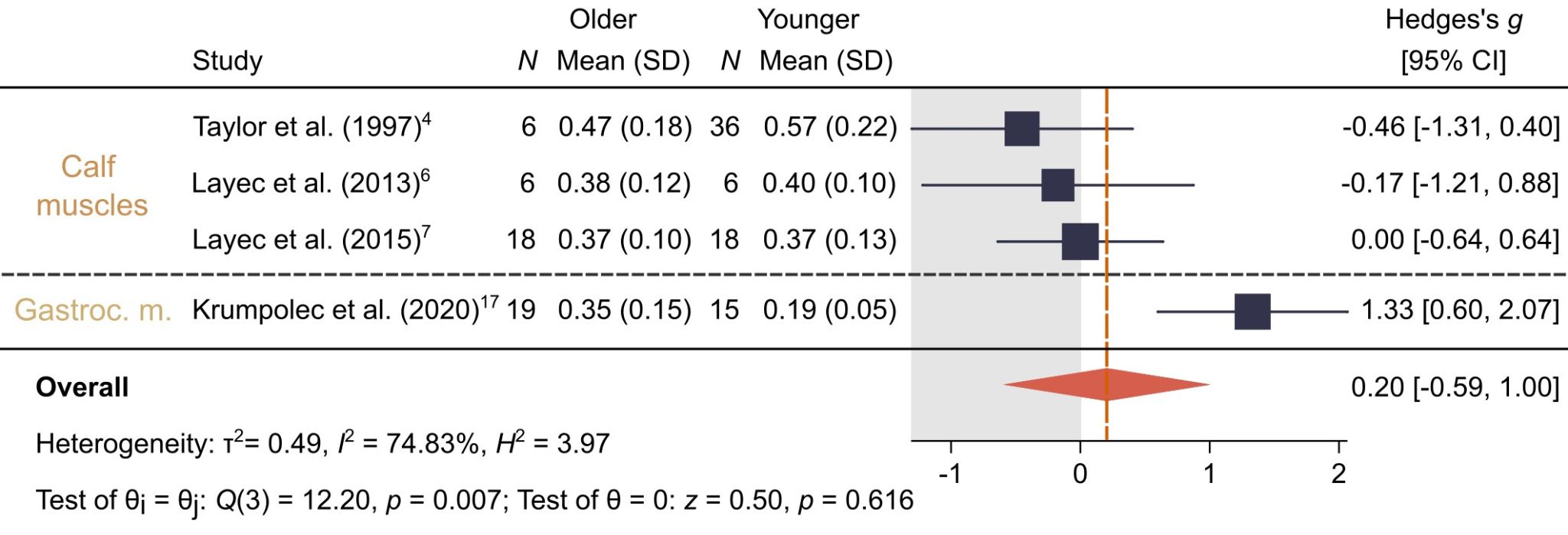

**Figure S5.** *Forest plot of the difference in the initial rate of phosphocreatine (PCr) recovery (Vi_PCr_) in muscles of the upper and lower leg between older and younger adults. Results are based on a random effects meta-analysis model fit using restricted maximum likelihood. Gastroc. m. = gastrocnemius medialis*

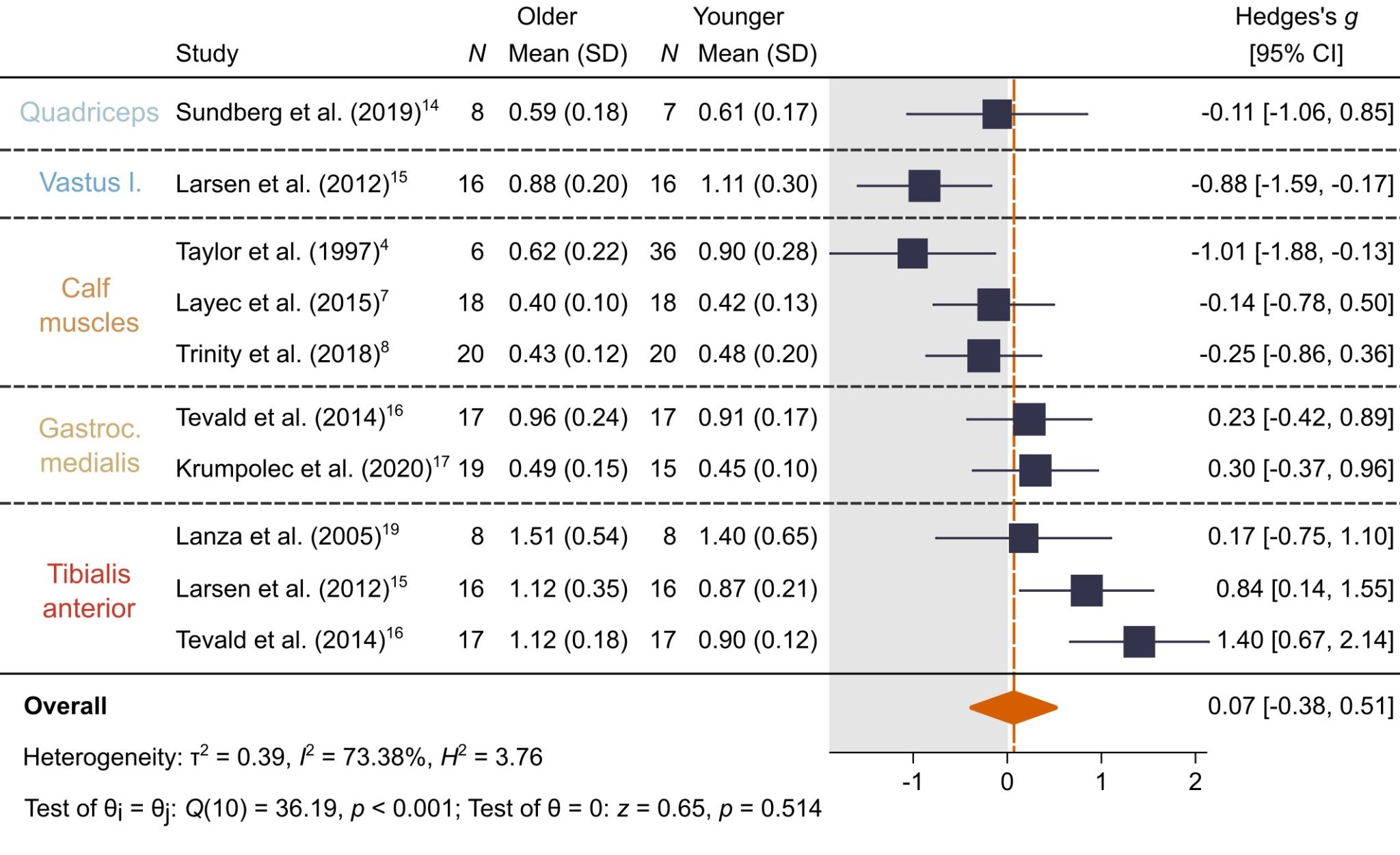

**Figure S6.** *Forest plot of the difference in the maximal rate of oxidative adenosine triphosphate (ATP) synthesis (Q_max_) in muscles of the upper and lower leg between older and younger adults. Results are based on a random effects meta-analysis model fit using restricted maximum likelihood. Gastroc. medialis = gastrocnemius medialis, Vastus l. = vastus lateralis*

**Supplementary Methods S1 - Search strategy**

**MEDLINE/EMBASE**

Search terms:

1. (exp Phosphorus AND exp Magnetic Resonance Spectroscopy);

2. ("31P magnetic resonance spectroscopy" OR ("31" adj1 (P or phosphorus) adj3 ("magnetic resonance spectroscopy" OR MRS OR NMR OR "nuclear magnetic resonance"))).ti,ab,kf.;

3. ((P OR phosphorus) adj3 ("magnetic resonance spectroscopy" OR MRS OR NMR OR "nuclear magnetic resonance")).ti,ab,kf.;

4. OR/1-3;

5. exp Muscle, Skeletal/ OR exp Muscular Atrophy/ OR exp Aging/ OR exp Geriatrics/ OR exp Sarcopenia/ OR Frail Elderly/ OR exp Geriatric Assessment/ OR exp Frailty/ OR exp "Aged, 80 and over"/ OR exp Middle Aged/ OR exp Adult/;

6. (musculoskelet* OR ((skelet* OR striat* OR "cross stripe*" OR trunk* OR voluntary OR atrophy OR age* OR aging OR frail* OR degener* OR shrink*OR weak* OR declin* OR deplet* OR diminish* OR insufficien* OR deteriorat* OR loss*) adj3 muscl*)).ti,ab,kf.;

7. (geriatric* OR adult* OR senior* OR age* OR old* OR frail* OR elder* OR senium or sarcopenia).ti,ab,kf.;

8. OR/5-7;

9. exp muscle contraction/ OR muscle fatigue/ OR exp muscle strength/ OR exp Oxygen Consumption/ OR exp Cell Respiration/ OR exp Mitochondrial Size/ OR exp Energy Metabolism/ OR exp Oxidative Stress/ OR exp Lactic Acid/ OR exp Phosphocreatine/ OR (exp Fatty Acids/ AND exp Oxidation-Reduction/) OR exp Oxidation-Reduction/ OR exp Adenosine Triphosphate/;

10. (muscl* adj3 (function* OR characteristic* OR strength* OR force* OR power* OR gain* OR fatigue OR assess* OR test* OR analy*)).ti,ab,kf.;

11. (bioenerg* OR oxidati* OR redox* OR (mitochondri* adj3 (metabolism OR size OR oxidat* OR respirat* OR stress OR clearance OR volume OR energ* OR bioenerg* OR redox OR reduct*))).ti,ab,kf.;

12. ((aerobic OR cell*) adj2 (oxidat* OR respirat* OR metaboli* OR redox OR reduction)).ti,ab,kf.;

13. ((creatine adj2 phosphate) OR (phosphocreatine OR phosphorylcreatine OR creatinephosphoric)).ti,ab,kf.;

14. (energ* adj2 (metaboli* OR cell* OR product* OR conver*)).ti,ab,kf.;

15. ((adenosine adj3 triphosphate) OR atp OR adenosinetriphosphate).ti,ab,kf.;

16. ((Intracell* OR cell* OR exercis* OR training) adj2 pH).ti,ab,kf.;

17. OR/9-16;

18. 4 AND 8 AND 17;

19. ((exp animals/ OR exp veterinary medicine/ OR animal*.jw.) NOT exp humans/) OR (animal* OR monkey* OR sheep OR ?ovine OR lamb* OR goat* OR pig* OR swine OR porcine OR pup* OR dog* OR canine OR bitch* OR beagle* OR feline OR rodent* OR rabbit* OR rat OR rats OR mouse OR mice OR murine).ti,kf.;

20. (magneti?ation adj2 transfer*).ti.;

21. (cancer* OR tumo?r OR oncolog*).ti.;

22. review.pt;

23. exp conference abstract/

24. OR/19-23

25. 18 NOT 24

**SPORTDiscus**

Search terms:

1. TX "31P magnetic resonance spectroscopy" OR TX ( ("31" adj1 (P or phosphorus) N3 ("magnetic resonance spectroscopy" or MRS or NMR or "nuclear magnetic resonance")) ) OR TX ( ((P or phosphorus) N3 ("magnetic resonance spectroscopy" or MRS or NMR or "nuclear magnetic resonance")) )

2. TX ((musculoskelet* or (muscl* N3 (skelet* or function* or characteristic* or strength* or force* or power* or fatigue or age* or aging) ) OR TX ( ((skelet* or striat* or "cross stripe*" or trunk* or voluntary or atrophy) N3 muscl*) ) OR TX ( (bioenerg* or oxidati* or (mitochondri* N3 (metabolism or oxidat* or respirat* or stress or clearance or volume or energ* or bioenerg*)) ))

3. S1 AND S2

**Cochrane Central Register of Controlled Trials (CENTRAL)**

Search terms:

#1. MeSH descriptor: [Magnetic Resonance Spectroscopy] explode all trees

#2. MeSH descriptor: [Phosphorus] explode all trees

#3. #1 AND #2

#4. ("31P magnetic resonance spectroscopy" or ("31" N1 (P or phosphorus) N3 ("magnetic resonance spectroscopy" or MRS or NMR or "nuclear magnetic resonance")))

#5. {OR #3 AND #4}

#6. (musculoskelet* or ((skelet* or striat* or "cross stripe*" or trunk* or voluntary or atrophy or age* or aging) NEAR/3 muscl*))

#7. #5 AND #6

**Supplementary Methods S2 - Adapted NOS questionnaire**
**Adapted Newcastle Ottawa Scale (NOS) – total of 11 stars to be awarded**

**1. Selection (max 7 stars)**

- **Representativeness of the sample (2 stars)**
  - Truly representative of the average population (2 star)
  - Somewhat representative (1 star) – i.e. recruited from the community setting
  - Selected group of users (e.g., volunteers, clinic patients) – i.e. recruited from the university or sports clubs – or recruitment strategy not reported (0 stars)
- **Sample size (1 star)**
  - Justified & satisfactory (1 star) - i.e. power calculation reported
  - Not justified (0 stars)
- **Non-respondents (1 star)**
  - Comparability between respondents & non-respondents described (1 star) – mentioned drop-outs or not
  - Unsatisfactory response rate / no description (0 stars)
- **Ascertainment of exposure (risk factor) (2 stars)**
  - Secure record (e.g., medical records, registry) (2 stars) – Physical activity logging using accelerometry or a combination of accelerometry and questionnaires
  - Structured interview/questionnaire (1 star) – Physical Activity monitoring using questionnaires only
  - Self-report / not validated (0 stars) – Not reported and or not measured

**2. Comparability (max 2 stars)**

- **Control for confounding factors (2 stars)**
  - Controlled for the most important factor (1 star) – Physical activity levels
  - Controlled for additional factor(s) (1 star) – gender, comorbidities, or other factors

**3. Outcome (max 3 stars)**

- **Assessment of outcome (1 star)**
  - 31P-MRS was used to measure tau PCr using a harmonised measurement protocol (1 star)
  - Non-validated measurement but described (0 stars)
- **Statistical test (2 stars)**
  - Clearly described, appropriate, and assumptions addressed (2 stars)
  - Clearly described, appropriate but assumptions not addressed (1 star)
  - Not appropriate or not described (0 stars)

**Supplementary Results S1**

|  | **Selection** | | | | **Comparability** | **Outcome** | | **Total** |
| --- | --- | --- | --- | --- | --- | --- | --- | --- |
| **Study** | 1 | 2 | 3 | 4 | 1 | 1 | 2 | max. 11 |
| Taylor et al. 1984^1^ |  |  |  |  |  |  | * | 1 |
| Distefano et al. 2018^13^ |  |  |  | ** | ** | * | * | 6 |
| Sundberg et al. 2019^14^ |  |  |  | ** | ** | * | ** | 7 |
| Larsen et al. 2012^15^ | * |  |  | ** | ** | * | * | 7 |
| Mancilla et al. 2023^2^ | * |  |  | * | ** | * | * | 6 |
| Bartlett et al. 2024^3^ |  |  |  | ** | ** | * | ** | 7 |
| Taylor et al. 1997^4^ |  |  |  |  | * | * | * | 3 |
| Wray et al. 2009^5^ |  |  |  | * | * | * | * | 4 |
| Layec et al. 2013^6^ |  |  |  | * | * | * | * | 4 |
| Layec et al. 2015^7^ | * |  |  | ** | * | * | * | 6 |
| Trinity et al. 2018^8^ |  |  |  | ** | ** | * | * | 6 |
| Lewsey et al. 2020^9^ | * |  |  | * | * | * | ** | 6 |
| Chilibeck et al. 1998^10^ |  |  |  |  |  | * | * | 2 |
| Waters et al. 2003^11^ | * |  |  | * | ** | * | * | 6 |
| Tevald et al. 2014^16^ | * |  |  | * | ** | * | * | 6 |
| Hart et al. 2015^12^ |  |  |  | ** | ** | * | * | 6 |
| Krumpolec et al. 2020^17^ | * |  |  | * | ** | * | * | 6 |
| Kent-Braun & Ng, 2000^18^ | * |  | * | ** | ** | * | * | 8 |
| Lanza et al. 2005^19^ | * |  |  | ** | * | * | * | 6 |
| Christie et al. 2014^20^ | * |  |  | ** | ** | * | ** | 8 |
